# Head-Direction drift in rat pups is consistent with an angular path-integration process

**DOI:** 10.1101/212852

**Authors:** Gilad Tocker, Eli Borodach, Tale L. Bjerknes, May-Britt Moser, Edvard I. Moser, Dori Derdikman

## Abstract

The sense of direction is a vital computation, whose neural basis is considered to be carried out by head-direction cells. One way to estimate head-direction is by integrating head angular-velocity over time. However, this process results in error accumulation resembling a random walk, proportional to 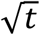, which constitutes a mark for a path integration process. In the present study we analyzed previously recorded data to quantify the drift in head-direction cells of rat pups before and after eye-opening. We found that in rat pups before eye-opening the drift propagated as a random walk, while in rats after eye-opening the drift was lower. This suggests that a path-integration process underlies the estimation of head-direction, such that before eye-opening the head-direction system runs in an open-loop manner and accumulates error. After eye-opening, visual-input, such as arena shape, helps to correct errors and thus compute the sense of direction accurately.

## Introduction

The sense of orientation is a vital computation, widespread in many animal species (Muller & Wehner 1988; Seelig & Jayaraman 2015). In mammals, the neural basis underlying orientation is considered to be carried out by head direction cells, which are cells that fire when the animal’s head points to a specific direction (Robertson et al. 1999; Finkelstein et al. 2016; Taube et al. 1990a; Taube et al. 1990b; Ranck 1985). The lack of specific sensors for head direction in mammals suggests that the brain infers the animal’s head direction indirectly from various sensory inputs. Previous studies showed that vestibular inputs are necessary for the formation of the head direction signal while visual inputs are important for anchoring the head direction signal to the external reference frame (Bjerknes et al. 2015; Tan et al. 2015; Muir et al. 2009; Valerio & Taube 2016; Taube et al. 1990b; Goodridge et al. 1998; Stackman & Taube 1997). However, the algorithm used by the brain to compute the animal’s head direction is unknown. One possible way to compute the head direction signal is by integrating the head’s angular velocity over time (Zhang 1996). If the noise source in such an integration process is uncorrelated and stationary, this process will result in accumulation of an error proportional to 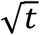 (because of its resemblance to Brownian motion). Therefore, a head direction drift proportional to 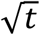 would be a mark for an angular path integration algorithm underlying the head direction system. Yet, it is difficult to measure the drift in adult rats because their head directionality remains stable for long periods of time even in the dark (Chen et al. 2016; Goodridge et al. 1998). Bjerknes et al. (2015) found that head-direction cells exist without visual input in rat pups before eye opening already at P13, but that their preferred directional tuning is unstable. This instability makes them a natural model for studying the drift of the head direction system.

In the present study we aimed to quantify the cell’s head direction drift in rat pups before and after eye opening. We found that in rats before eye opening accumulation of drift resembled the power law expected out of an angular path integration process. In contrast, in rats after eye opening, the accumulation of drift was lower than expected.

We furthermore checked how environmental shape affected head direction drift. A square environment has a 4-fold rotational symmetry, while a circular environment has an infinite-fold rotational symmetry. Therefore, a square environment bares more information about the rat’s head direction. We found that the magnitude of the drift was dependent on the geometrical shape of the environment. Together, our findings support a vestibular-dependent angular path-integration algorithm, whose error accumulates over time, which can be corrected by vision. Furthermore, the rat uses the environment’s geometrical shape to correct this accumulated error.

## Material and Methods

### Subjects

For the analysis of rat pups we used previously published data from Bjerknes et al. (2015). A total of 163 cells were recorded from 14 rats. 131 of these cells were recorded twice. 86 cells were recorded in a circular arena during the last 3-4 days before eye-opening (P11-P13). 62 of these cells were recorded twice. Recordings were done on P11 in one rat, P12 in three rats, P13 in six rats, P14 in eight rats, and P15 in one rat. 52 cells were recorded in a circular environment 1-2 days after eye-opening. 49 of these cells were recorded twice. Recordings were done on P14 in one rat, P15 in eight rats and P16 in 6 rats. 25 cells were recorded in a square environment 1-2 days after eye-opening. 20 of these cells were recoded twice. Recordings in a square environment were done on P15 in two rats and on P16 in two rats. The tetrodes were placed in presubiculum in seven rats, in parasubiculum in four rats, at the border between pre- and parasubiculum in two rats, and in medial entorhinal cortex (MEC) in one rat. The tetrodes were distributed across deep and superficial layers of pre- and parasubiculum and deep layers of MEC. The pups moved freely across the recording arena and covered the entire range of head directions. For the analysis of the adult groups we used previously published data from (Sargolini et al. 2006; Bonnevie et al. 2013; Derdikman et al. 2009). A total of 408 head-direction cells were analyzed. All cells were recorded from the MEC. 93 of these cells were recorded in a circular environment. 315cells were recorded in a square environment.

### Head Directionality

Head direction tuning curves were generated by dividing the number of spikes fired when the rat faced a particular direction (in bins of 6°) by the total amount of time the rat spent facing that direction. The resulting tuning curve was then smoothed using a 10-point hamming window. The strength of head directionality was estimated by computing the length of the mean vector (Rayleigh vector) for the circular distribution of the tuning curve.

### Classification of head direction cells

Classification of head direction cells was done by determining the significance of the cell’s head directionality, using a shuffling procedure. The null hypothesis was that the head directionality of the cell was not caused by a dependence of the cell’s firing rate on the rat’s head direction during the session. Therefore, for each cell, the whole time series of spikes was shifted in time by a random (uniformly-distributed) interval between 0 and the time length of the session (600 seconds). Spikes which were out-bounded were rotated cyclically to the beginning. This procedure preserved the temporal structure of the cell’s spiking activity and the rat’s head direction behavior during the session, but dissociated between the two. We repeated this process 100 times for each cell. If the Rayleigh vector of a cell was longer than the 95^th^ percentile of the shuffled cells Rayleigh vectors distribution, the null hypothesis was rejected, meaning that the head directionality of the cell was most likely due to the cell firing at a specific head direction of the rat. Cells that passed this significance threshold were classified as head direction cells in our study.

### Quantifying the cell’s drift

To quantify the cell’s drift we first calculated for each pair of spikes in the spike train, (1) the time interval between the spikes (ΔT) and, (2) the head direction difference between the spikes (Δ*HD*). Then, we partitioned the log(Δ*Τ*) log(Δ*HD*) space into bins of 0.1×0.1 log(sec)log(degrees) and counted the number of spike-pairs in each bin. We subsequently smoothed the histogram using a Gaussian kernel with a standard deviation of *σ* =1.5 cm. Finally, we divided the number of spike-pairs in each bin by the total amount of spikes pair-sharing the same log(Δ*Τ*)-reading, to create a probability function of log(ΔHD) given log(Δ*Τ*). We then fit a regression line (slope and intersection) between all log (ΔT) values and the most probable log (ΔHD) values that corresponded to them.

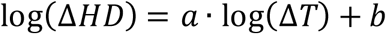

where *a* is the slope and *b* is the intersection. We ignored ΔT smaller than 2 sec.

### Evaluating the drift significance of population of cells

To evaluate the significance of the drift across the whole cell population, a shuffling procedure was used. The null hypothesis was that the temporal structure of the cell’s spiking activity was independent from the rat’s behavior. Therefore, the whole series of spikes was shifted in time by a random (uniformly-distributed) interval between 0 seconds and the length of a single session (600 seconds). Spikes which were out-bounded were rotated cyclically to the beginning. For each shuffled cell, we calculated the cell’s drift in a similar manner to the calculation used for the real cell, and then we fitted a regression line (slope and intersection) similar to that described in the previous section. We repeated this process 1000 times for each cell. The median slope and intersection of the original population was then compared to the distributions of median slopes and median intersections of the shuffled populations.

### Simulating cells

To simulate head direction cells with no drift, we simulated non-homogeneous Poisson cells. At each millisecond t, λ(*t*) was chosen as the value of the bin in a Gaussian head direction tuning curve that was closest to the instantaneous head direction of the rat. The Gaussian head direction tuning curve function was:

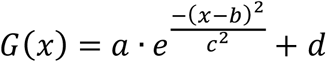

Parameters a=15, c=50 and d=0.2 were selected by fitting a Gaussian function to a representative head direction cell of an adult rat with no drift (results did not changed significantly after fitting Gaussian to a different cell). Parameter b which is the preferred head direction of the cell was selected randomly from a uniform distribution between 0 and 360.

To simulate drifting head direction cells, we simulated the head directionality as in the simulation of head direction cells with no drift. The insertion of drift was done by shifting the Gaussian head direction tuning curve at each time step (1 millisecond) randomly by ε degrees either to the left or to the right. Values of the Gaussian distribution that drifted beyond 360° were wrapped cyclically to the beginning. The simulation of the non-cyclic drift was done by allowing the Gaussian head direction tuning curve to drift beyond the 360° bound. λ(*t*) was chosen as the bin in the Gaussian distribution that was closest to the instantaneous head direction of the rat, modulo 360. Note that the cells were drifting continuously during the simulation and there was no correction mechanism.

Choosing ε values: we tested several different values of ε (0, 0.2, 0.4, 0.6, 0.8, 1). ε=0 is a simulation of head direction with no drift. We chose ε=1 to be our drift upper limit because in this case 87% of the cells do not show significant head directionality (Figure 1E, F). Meaning that the cells lose their head directionality in this levels of drift. If this was the case in the real recorded data we could not have detected these head direction cells. However, in the real data we did find head direction cells (51% of all recorded cells passed the head directionality criteria).

**Figure 1.**
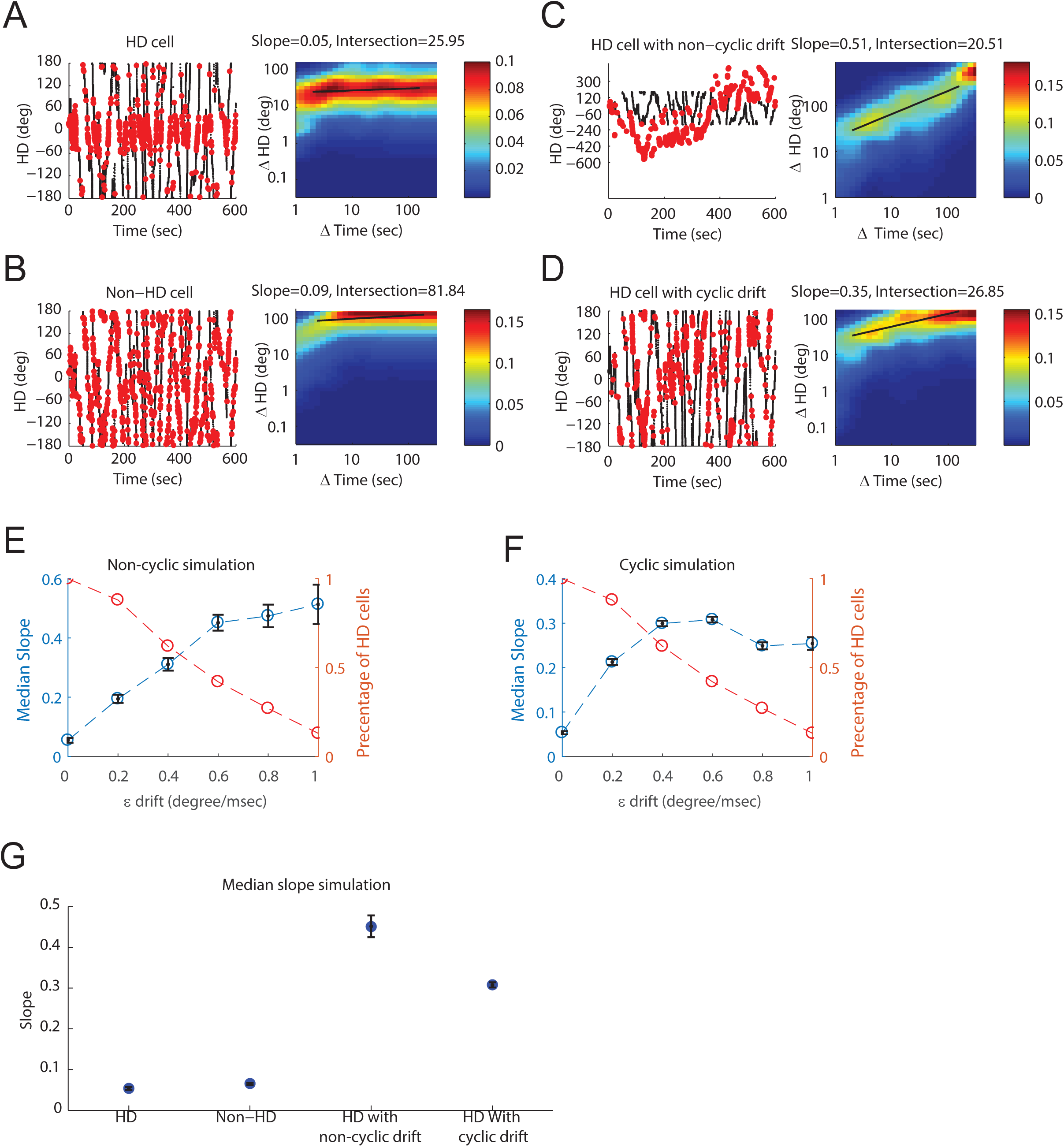
Drift quantification results of simulated cells. **A-D** are examples of simulated cells. Left column, in black is the head direction of the rat during the session. The red dots mark the rat’s head direction while the cell fired spikes. Right column, is the probability function of log(ΔHD) given log(Δ*Τ*). The black line is the regression line (slope and intersection) fitted to a**l** log (ΔT) values and the most probable log (ΔHD) values that correspond to them. **A**, an example of a simulated head direction cell with no drift. **B**, an example of a simulated non-head direction cell. **C**, an example of a simulated head direction with non-cyclic drift. **D**, an example of a head direction with cyclic drift. **E**, simulation of drifting population of cells in the cyclic method, for different drift values (degree/msec). x axis *ε*– drift values, Left y axis is the median slope of the cells, Right y axis is the percentage of significant HD cells. **F**, simulation of drifting population of cells in the non-cyclic method for different drift values (degree/msec). x axis – drift values, Left y axis is the median slope of the cells, Right y axis is the percentage of significant HD cells. **G**, simulated population results. In blue are the median regression line slopes of all simulated cells (we used a drift value of *ε* =0.6 (degree/msec)).

To simulate non-head directions cells, we simulated homogeneous Poisson cells. λ was chosen to be 4. The results did not change significantly when using a different λ(2, 3, 6 – data not shown).

For all simulations, we used a trajectory of a rat pup at P14 running in a circular environment.

## Results

### Simulation of cells

We started investigating the drift of head direction cells, by first asking what would be the expected head direction drift of 3 different kinds of cells: (1) head direction cells with no drift (Figure 1A) (2) non-head direction cells, meaning Poisson ce**l** s with constant λ along the session (Figure 1B), and (3) head direction cells with drift (Figure 1C, D). We simulated 100 cells of each kind and quantified their drift. Accumulated noise proportional to 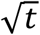 is the theoretical characteristic of an iterative process with zero mean noise, which should behave like Brownian motion. However, the cyclic nature of head direction variable can cause an underestimation of this drift. To address this issue, we simulated the head directionality drift of cells by two different methods. In the first method, we used head direction as a non-cyclic variable, meaning that the drifted head direction could grow beyond 360 degrees (Figure 1C). In the second method, we used head direction as a cyclic variable between 0 and 360 degrees as in the real life data (Figure 1D). Note that in both methods the head direction cells were drifting continuously without any correction mechanism (see Material and Methods). Quantification of drift was done by first computing (1) the time interval between a**l** spike pairs of the ce**l** (ΔT), and (2) the head direction difference between these spike pairs (Δ*HD*), second, normalizing the spike pairs into conditional probability of log Δ*HD* given log ΔT, and lastly fitting a regression line between log ΔT and the most probable log Δ*HD* (see Materials and Methods).

### Drift quantification results of simulated cells

We found that the median log-drift slope of the simulated head direction cells with no drift was close to zero: 0.0534 ± 0.03 as expected (Figure 1A, G). The log-drift slope of the simulated Poisson non-head direction cells was 0.065 ± 0.002. The small log-slope of the drift was a result of the Poisson nature of these cells (Figure 1B, G). The median intersection of non-head direction cells was significantly higher than the median intersection of head direction cells (88 ± 1, 37 ± 1, p<10^−34^; Wilcoxon Ranksum test, Figure S1A).

The median log-drift slope of drifting cells in the non-cyclic simulation method was close to 0.5 for drift higher than 0.6 degrees per msec (0.4515± 69, 0.4744 ± 0.0381, and 0.51 ± 0.0663 for drifts 0.6, 0.8, and 1 degree per msec) (Figure 1E) as expected from a diffusion-like process (Figure 1C, G). In contrast, the median log-drift slope in the cyclic simulation method was lower (0.3079 ±0.0065 was the highest for 0.6 degrees per msec drift) (Figure 1D, E, G). The lower values of the cyclic case was a result of the cyclic variable tending to reduce large values of drift. Thus, we expect head direction drift originating from an iterative processes to have a maximum error accumulation slope of about ∼0.3.

### Drift quantification results of head directions cells in rat pups

After we simulated a population of cells and found the expected drift of each kind of cell, as described above, we continued and applied the same methods to real recorded head direction cells, and asked how their drift meets our predictions from simulation. We analyzed 294 previously recorded cells from Bjerknes et al. (2015). Cells were recorded from the pre- and para- subiculum of 14 rat pups while they moved freely in the arena. Pre-eye-opening cells were recorded on P11-P14. Post-eye-opening cells were recorded on P15-P16. Cells were classified as head direction cells if their Rayleigh vector length was longer than the 95^th^ percentile of a shuffled distribution. 152 out of 294 cells passed this criterion and therefore were defined as head direction cells and were included in further analysis.

The head directionality of many of the cells in rat pups drifted over time (Figure 2B-E). We thus aimed at measuring the gradual accumulation of drift, in rats before eye-opening vs. after eye-opening. We analyzed 56 cells recorded in rat pups before eye-opening (P11-P14) and 60 cells recorded in rat pups after eye opening (P15-P16). All cells were recorded while rat pups were foraging for food in a circular environment. The median log-slope of cells recorded from rat pups before eye opening was 0.3098 ± 0.0265, significantly higher (p<10^−4^; Wilcoxon Ranksum test) than the drift in rat pups after eye opening, which had a log-slope of 0.1256 ± 0.0238 (Figure 2F). In adult rats the slope was small (log-slope=0.0774 ± 0.0101) (Figure 2F). There was no significant difference between the drift log-slope in rat pups before eye-opening and the maximum drift log-slope of simulated drifting cells (*ε* =0.6 degree per msec drift, p=0.9228; Wilcoxon Ranksum test) (Figure 2G). In contrast, the drift log-slope after eye-opening was significantly lower than the drift log-slope of simulated drifting cells (Figure 2G, p<10^−5^; Wilcoxon Ranksum test).

**Figure 2.**
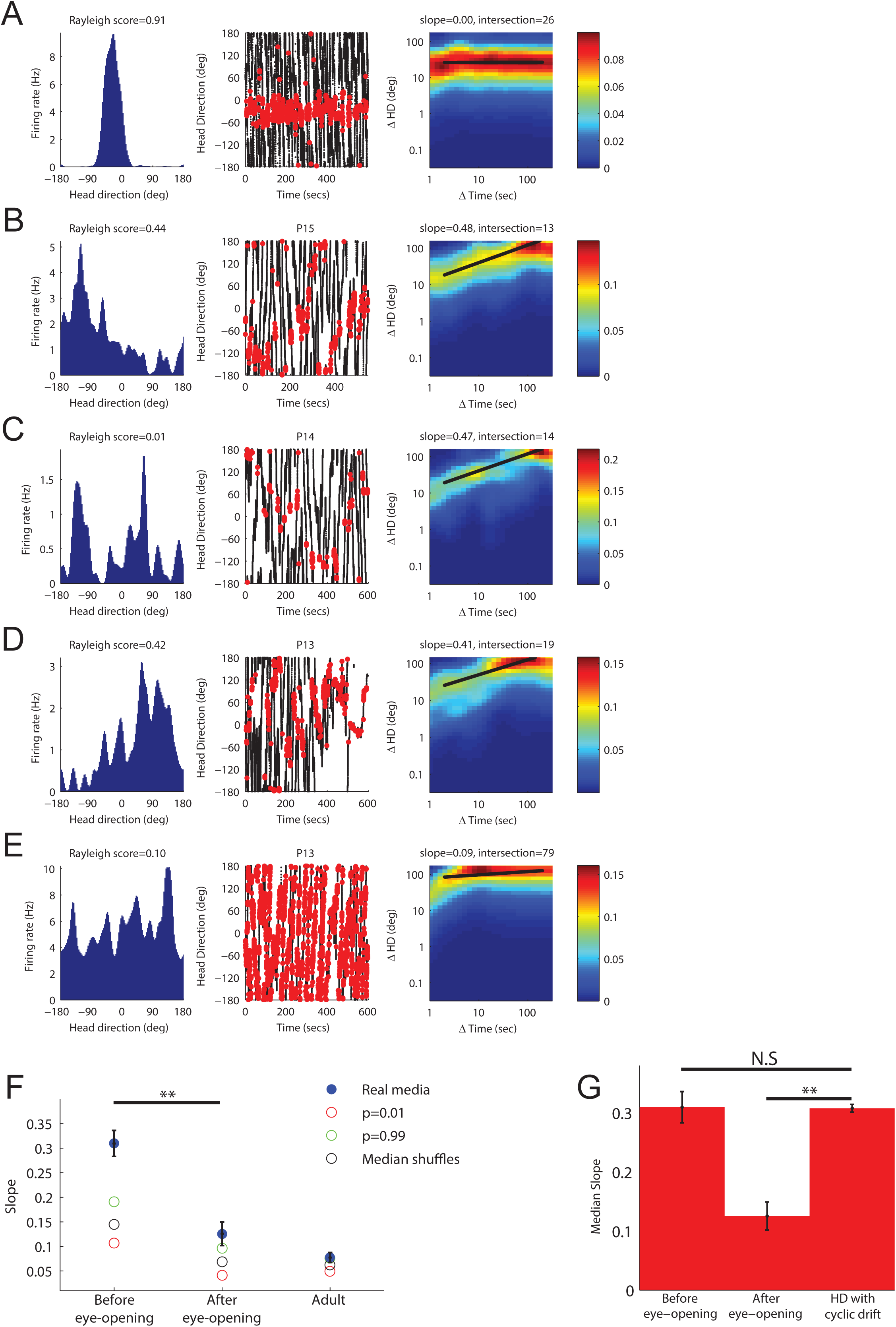
Drift quantification results of head direction cells in rat pups. **A-E** Examples of 5 recorded cells. Left column: the head direction tuning curve of the cell. Middle column: in black is the head direction of the rat during the session. The red dots mark the rat’s head direction while the cell fired spikes. Right column: the probability distribution of log(ΔHD) given log(Δ*Τ*). The black line is the regression line (slope and intersection) fitted to a**l** log (ΔT) values and the most probable log (ΔHD) values that correspond to them. **A**, example of a head direction cell in an adult rat. **B-D**, examples of drifting head direction cells in rat pups. **E**, example of a non-head direction cell. **F**, Quantification of drift in rats before eye-opening, after eye-opening and in adults. In blue are the median regression line slopes of cell populations recorded from rats before eye opening, after eye-opening and from adults. In red, black and green are the 1^st^ median and 99^th^ percentile of a shuffle distribution, respectively. **G**, drift in rats before eye-opening, after eye-opening and simulation of cyclic drift (drift value used -*ε* =0.6 (degree/msec)). N.S – not significant.

We applied a shuffling procedure on all cells to check if the drift we found was merely an artifact of the rat’s behavior and of the independent spiking activity. The median log-drift slope of cells of rats before eye-opening and after eye-opening was significantly higher than of the shuffled distribution (p<0.01) (Figure 2F), suggesting that the rat’s behavior alone could not account for the drift.

### Environmental geometry affects the cell’s drift

We next asked whether the environment’s geometry affects the cells’ drift. We analyzed a population of 35 cells recorded from rats-pups after eye opening foraging for food in a square environment together with 315 cells from adult rats foraging for food in a square environment, and compared them to 60 cells that were recorded in rats-pups after eye-opening foraging for food in circular environment together with 93 adult rats foraging for food in circular environment. We found that the drift, when rats ran in a square environment, was significantly lower (slope=0.0271 ± 0.0065) than the drift when rats ran in a circular environment (Figure 3) (slope= 0.0857 ± 0.0117, p<10^−9^; Wilcoxon Ranksum test).

**Figure 3.**
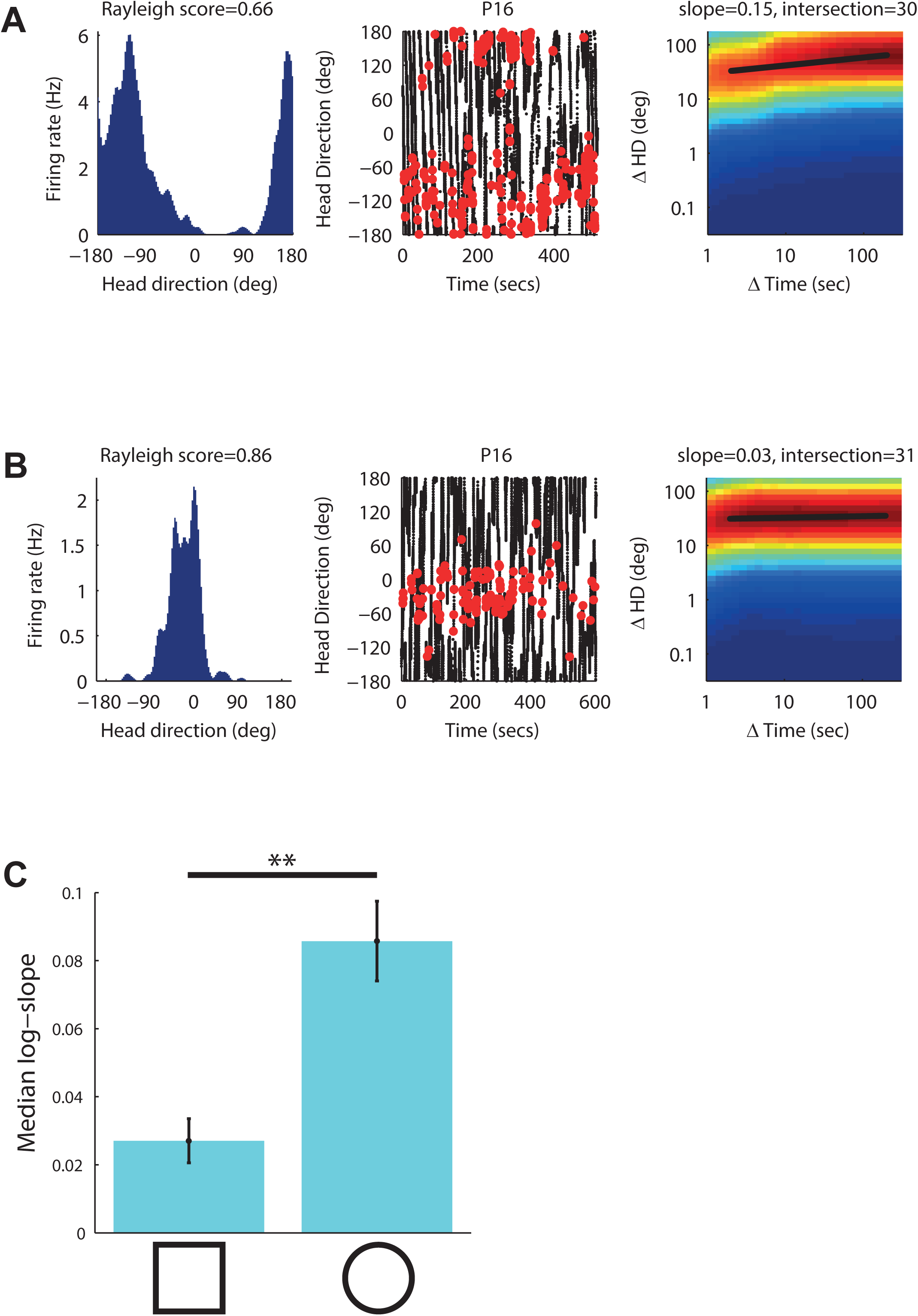
Environmental geometry affects the cell’s drift. **A**, Example for a cell recorded from a rat running in a circular arena. **B**, Example for a cell recorded from rat running in a square arena. Left column: the head direction tuning curve of the cell. Middle column: in black is the head direction of the rat during the session; red dots mark the rat’s head direction while the cell fired spikes. Right column: the probability function of log(ΔHD) given log(Δ*Τ*). The black line is the regression line (slope and intersection) fitted to all log (ΔT) values and the most probable log (ΔHD) values that correspond to them. **C**, The median log-slope of cells recorded from rats running in a square environment and the median log-slope of cells recorded from rats running in a circular environment.

## Discussion

In the present study we found that the head direction drift in rats before eye opening (P13-14) followed a power law of *∼t*^0.31^, while in rats after eye opening (P15-16) it followed a power law of ∼*t*^0.12^. The accumulation of drift in rats before eye opening resembled the power law expected out of an angular path integration process, and in rats after eye opening was lower than expected. In rats after eye opening and adults the drift in a square environment was smaller than in a circular environment.

Many theoretical studies proposed that the head direction signal was computed by an angular path integration process, therefore drifting and accumulating error over time (Skaggs et al. 1995; Redish et al. 1996; Zhang 1996). However, Although experimental work has shown that the preferred tuning direction of head direction cells changed when the rat was devoid of visual input (Bjerknes et al. 2015; Goodridge et al. 1998), these studies did not directly quantify the nature of the drift. By taking advantage of the head direction drift in rat pups before eye-opening, we were able to quantify the drift in this system. This quantification has implications for the algorithm underlying the estimation of head direction. The fact that the drift power law we found in rats before eye-opening was not significantly different from the expected drift in a path integration process (Figure 2G), suggests that an integration process underlies the estimation of head direction. Moreover, this was the upper limit of drift without any correction mechanism, which suggests that in rat pups before eye-opening there is almost no correction mechanism, and the system runs in practice in an open loop manner. The fact that this integration process occurs without the visual input suggests that a different source of input is being integrated, such as vestibular input. This extends adult studies showing the contribution of the vestibular system to the formation of head direction signal (Valerio & Taube 2016) by unraveling the nature of this vestibular contribution. The decrease in the drift after eye-opening implies an error correction mechanism that depends on the visual input.

Furthermore, a previous study suggested that the environment boundaries could be used as a mechanism for correcting the error accumulated in the grid cells code (Hardcastle et al. 2015). Our study suggests that the head direction code uses the information embedded in the geometrical shape of the environment to correct the accumulated error. The source of this error correction could come directly from salient geometric features in the environment such as walls and corners, or, from the border cells population that fire in an adult-like manner from the first days of exposure to an open environment (Bjerknes et al. 2014). Our data-set lacked a population of cells recorded in a square environment before eye-opening and so could not address the question whether the source of this correcting signal was visual or tactile. Future studies should address this question.

Computing the sense of direction instantaneously and accurately is vital for survival of humans and animals. Together, our findings support a vestibular path-integration algorithm that gives instantaneous sense of direction, but accumulates error over time. In addition, the environmental geometry, among other cues, helps to correct this accumulated error, and thus helps to compute the sense of direction accurately. We thus show here for the first time that the drift of the head-direction cells before eye-opening behaves like a Brownian-motion diffusion process. More generally, we provide here an example of how measuring the noise in a brain process, and not in the signal itself, can shed light on the computation performed by the brain.

---

**Figure S1.**
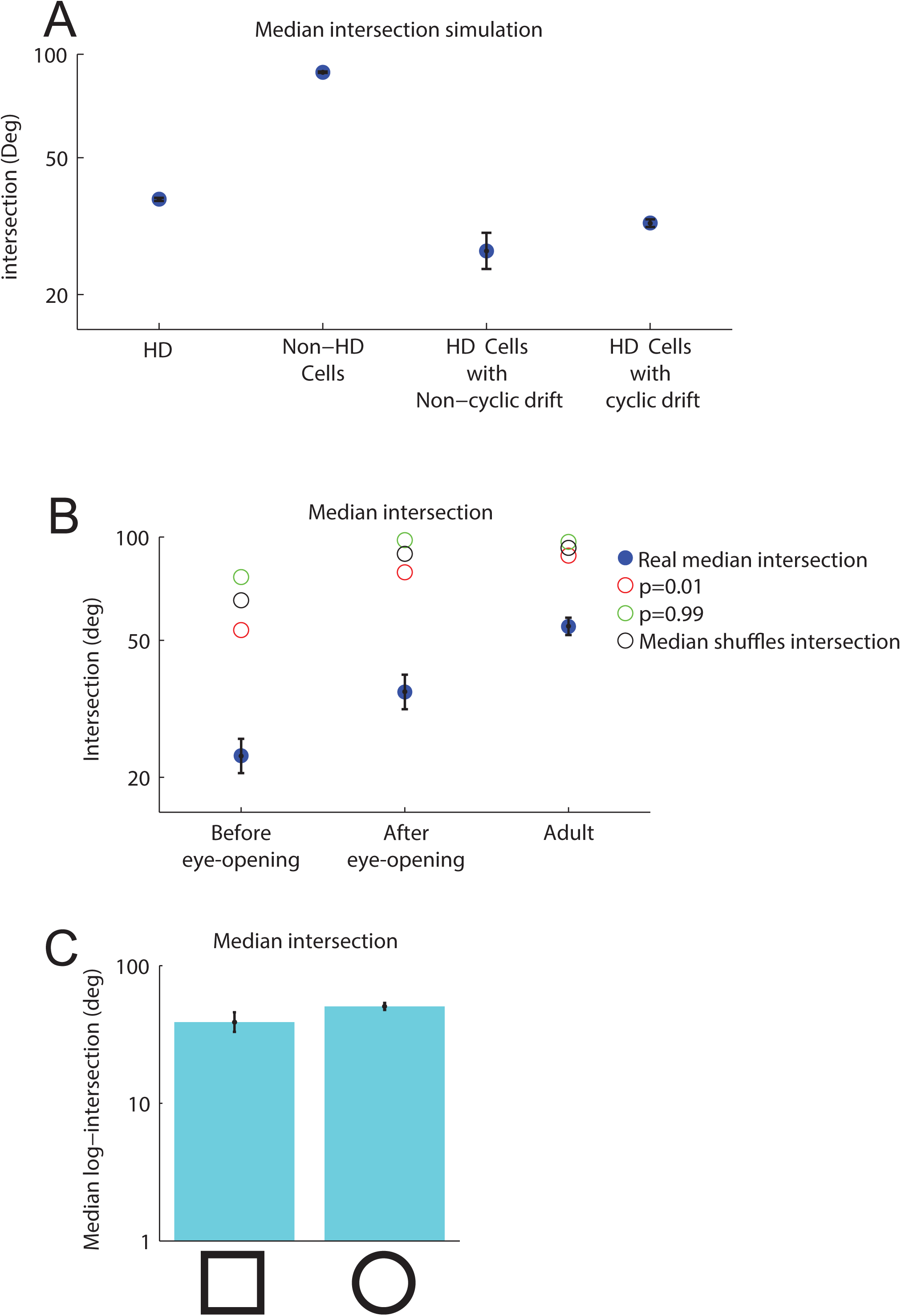
Quantification of intersection parameter. **A**, Simulated population results. Blue: the median regression line intersections of all simulated cells. **B**, Blue: the median regression line intersection of cell populations recorded from rats before eye opening, after eye-opening and adult. Red, black and green are the 1^st^ percentile, the median, and the 99^th^ percentile of the shuffle distribution, respectively. **C**, The median log-intersection of cells recorded from rats running in a square environment and the median log-intersection of cells recorded from rats running in a circular environment.

